# Computational prediction and analysis of very high risk single nucleotide polymorphisms in human cytochrome P450 oxidoreductase gene

**DOI:** 10.1101/590067

**Authors:** Dhas D Benet Bosco, K Rajalakshmi, S Suganya, P Pavani, K Yaswanth

## Abstract

Cytochrome P450 oxidoreductase (POR) is a highly polymorphic gene which is involved in metabolism of drugs and steroids through transfer of electron from NADPH to all CYP enzymes. In this study, we attempt to identify the very high risk single nucleotide polymorphisms in POR gene that would affect phenotype of the enzyme. The genetic variants in POR gene were retrieved from databases and analyzed with appropriate online computation tools. Very high risk non-synonymous SNPs were identified with 12 different sequence and structure homology based tools and evolutionary conservation tool (Consurf). Further the phenotype effect of the variant was assessed with MutPred2 and LigPlot. The very high risk non-coding variants were predicted with HaploReg V4 and RegulomeDB tools. The very high risk SNPs that may affect miRNA target sites were screened using PolymiRTs v3.0, miRNA SNP v2.0 and MirSNP. Among 4,601 variants in POR gene, 58 missense variants, 8 non-coding variants and three SNPs in miRNA target sites were found to be very high risk. These very high risk variants may regulate the expression and activity of cytochrome P450 oxidoreductase enzyme leading to differential drug and steroid metabolism by CYP enzymes.

## INTRODUCTION

Cytochrome P450 oxidoreductase (POR) is a membrane bound enzyme, found abundant in endoplasmic reticulum and plays major role in steroid and drug metabolism by transferring electron from nicotinamide adenine dinucleotide phosphate (NADPH) to cytochrome P450 enzymes [1]. POR protein structure has three binding domains for NADPH, FAD, FMN and α-helical domain connecting FAD and FMN domains [2]. POR gene is highly polymorphic and is located in 7^th^ chromosome consisting of 15 protein-coding exons and one non-coding exon that helps in initiation of transcription [3]. Single nucleotide polymorphisms (SNPs) in POR gene was found to affect the expression and activity of POR enzyme, significantly [4,5,6,7]. A study by Huang et al., found that POR has 140 SNPs with more than 1% minor allele frequency (MAF) among four different populations in San Francisco, African-American, Caucasian-American, Mexican-American, Asian American, respectively [8]. The most commonly studied polymorphism is POR*28 (A503V) which has MAF of 19.1, 26.4, 37.5 and 40% among African-American, Caucasian, Chinese and Japanese, respectively [8,9]. Other polymorphisms like -173C>A and - 208C>T have higher MAF in American and European populations, however both were not prevalent among Chinese population [10]. These population based variations has to be explored to understand differential enzymatic activity of POR and corresponding CYP enzymes which are involved in drug metabolism. This study is attempted to predict the very high risk SNPs in POR gene that may exert critical consequences in the structure and function of POR protein which in turn will influence differential drug metabolism.

## MATERIAL AND METHODS

### Sequence retrieval

The SNPs in POR gene were retrieved using the Table Browser tool in UCSC Genome Browser (https://genome.ucsc.edu/) and verified with dbSNP (https://www.ncbi.nlm.nih.gov/snp), ENSEMBL (http://www.ensembl.org), 1000 Genomes (https://www.ncbi.nlm.nih.gov/variation/tools/1000genomes).

### Analysis of non synonymous SNPs

The functional effect of missense variants present in *POR* gene were annotated using UCSC Genome Browser tool, Variant Annotation Integrator (VAI) (https://genome.ucsc.edu/) which is based on the Database of Non-synonymous Functional Predictions (dbNSFP) that includes the prediction tools SIFT, PolyPhen2, MutationTaster, MutationAssessor and Likelihood ratio test (LRT). In addition to VAI, in order to improve prediction, seven other web server based tools such as PANTHER (www.pantherdb.org/tools/), Meta SNP (http://snps.biofold.org/meta-snp/), nsSNPAnalyzer (http://snpanalyzer.uthsc.edu/), SNAP2 (https://rostlab.org/services/snap2web/), PhD-SNP (http://snps.biofold.org/phd-snp/phd-snp.html), SNPs & GO (http://snps.path.uab.edu/snps-and-go/snps-and-go-3d.html) and PMut (http://mmb.pcb.ub.es/PMut/). The missense SNPs which were predicted as deleterious by more than ten tools are chosen as highly deleterious variants and analyzed with ConSurf server (http://consurf.tau.ac.il/) for their evolutionary conservation based on phylogenetic analysis. The conserved variants with score 8 and 9 were considered as very high risk nsSNPs.

### Structural and Functional consequences of very high risk nsSNPs

The very high risk nsSNPs were further analyzed for their structural and functional consequences using the tool, MutPred2 (http://mutpred.mutdb.org/index.html). MutPred2 predicts deleterious mutations based on their structural and functional effects on molecular mechanisms. This tool has been experimentally validated using yeast two-hybrid system to study the high MutPred2 scoring mutations associated with mendelian disorders. The variants with MutPred2 score > 0.5 were taken as pathogenic and the molecular mechanisms disrupted were found using a threshold p-value of 0.05. The very high risk variants present in the binding sites of POR protein for the ligands, flavin mononucleotide (FMN), flavin adenine dinucleotide (FAD) and nicotinamide adenine dinucleotide phosphate (NADP) were identified using LigPlot tool associated with PDBSum database (www.ebi.ac.uk/thornton-srv/databases/cgi-bin/pdbsum/GetPage.pl?pdbcode=index.html).

### Annotation of non-coding SNPs

The non-coding SNPs (intronic and UTR variants) retrieved from databases were screened based on their global minor allele frequency and those with more than one percentage are annotated for their regulatory actions using the tools, HaploReg v4.1 (http://archive.broadinstitute.org/mammals/haploreg/haploreg_v4.php) and RegulomeDB (http://www.regulomedb.org/).

### SNPs in miRNA target sites

The very high risk SNPs in the miRNA target sites were identified using the tools, PolymiRTs v3.0 (http://compbio.uthsc.edu/miRSNP/), miRNA SNP v2.0 (http://bioinfo.life.hust.edu.cn/miRNASNP2/) and MirSNP (http://bioinfo.bjmu.edu.cn/mirsnp/).

## RESULTS

Cytochrome P450 oxidoreductase (POR) is a membrane bound enzyme having 680 amino acids (UniProt ID: P16435 & Q59ED7) with a gene size of 71,754 bp (NC_000007.14) that includes 15 coding exons and one non-coding exon. The genetic variants in POR gene were retrieved from UCSC Genome Browser and cross verified with dbSNP and ENSEMBL (**Table S1**). POR gene was found to have 4,601 genetic variants including SNPs (93.5%), insertions (3.4%), deletions (2.9%) and others including in-dels and MNPs (Multi Nucleotide Polymorphisms) (Fig. 1A). Based on the function, the SNPs in POR consists of intronic (86.6%), non-synonymous (8.4%), coding synonymous (3.8%) and untranslated region (UTR) variants (1.2 %) (Fig. 1B). The distribution of non-synonymous variants in coding exons is schematically represented in Fig. 1C.

**Fig. 1:**
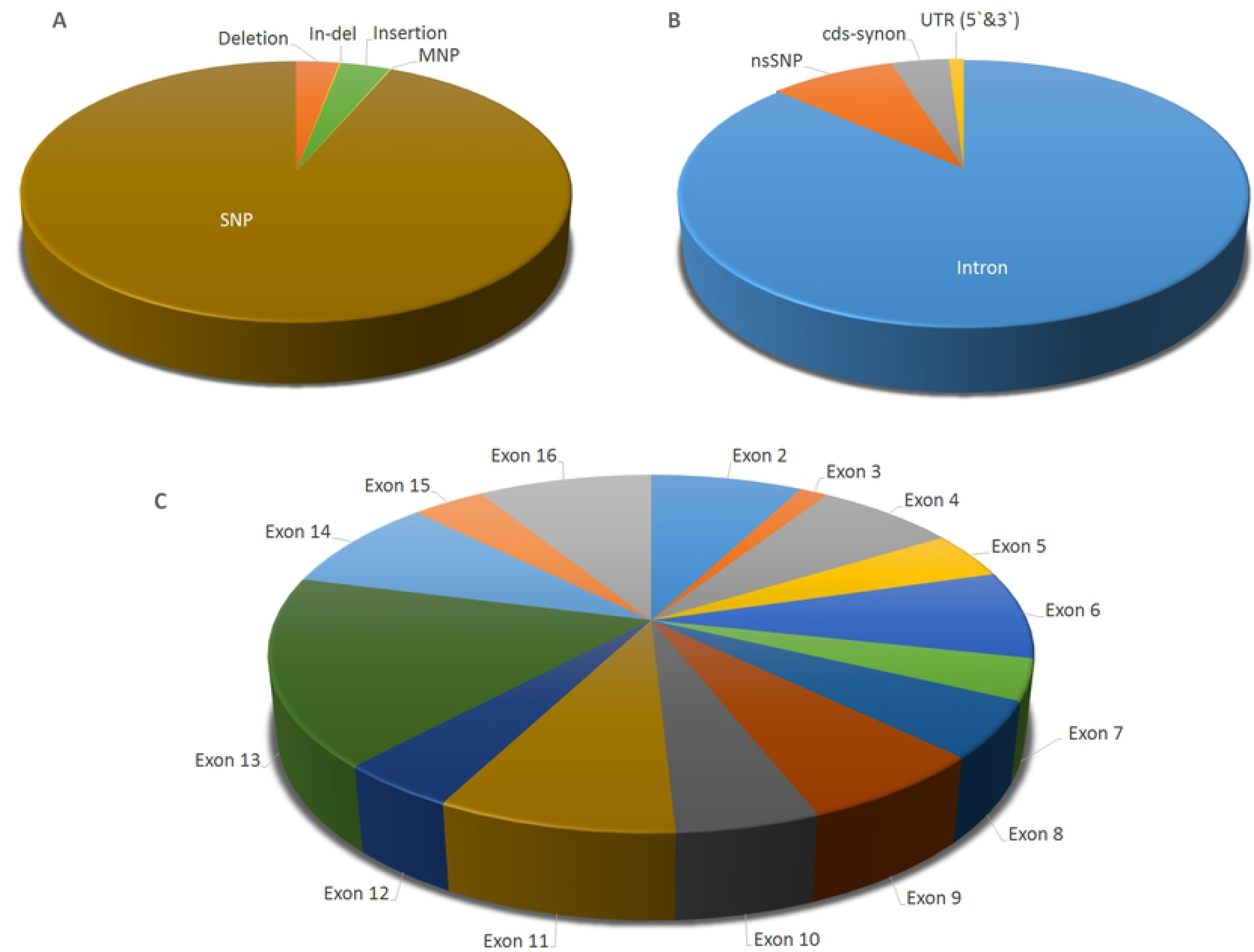
Distribution of POR gene variants based on: (A) class of variants (B) location in gene structure (C) coding nonsynonymous SNPs

POR gene has a total number of 376 nsSNPs that include 352 missense and 24 nonsense variants. Among the 352 missense variants, 23 SNPs can lead to more than one type of aminoacid substitution based on the codon change, for example, in rs201513102 the substitution of the ancestral allele G by A forms the codon ATG which changes Valine to Methionine, whereas substitution by C forms the codon CTG that introduces Leucine in place of Valine. Both substitutions were analyzed as distinct SNPs to identify their own biological significance. The missense variants were analyzed with 12 different prediction tools to identify the deleterious SNPs based on sequence and structural homology (**Table S2**). The prediction of each tool, benign or deleterious is represented in Fig.2A with green and red colors respectively, for each missense variant. Out of 352 missense variants, 31 SNPs by 11/12 tools and 56 SNPs by 12/12 tools were predicted as deleterious, whereas 15 SNPs were found to be benign by 12/12 tools (Fig.2B). The distribution of benign and deleterious variants in the exons are schematically described in Fig.2C. The color palette shows the number of tools that predicted a particular SNP as deleterious. Exon 13 was found to have high number of missense variants SNPs with 20 high risk nsSNPs (predicted by more than ten tools), in contrast, exons 2, 3 and 11 have no high risk nsSNPs. The commonly found variant, A503V was predicted to be deleterious only by PANTHER tool and hence it was not included in the high risk nSNPs list.

**Fig.2:**
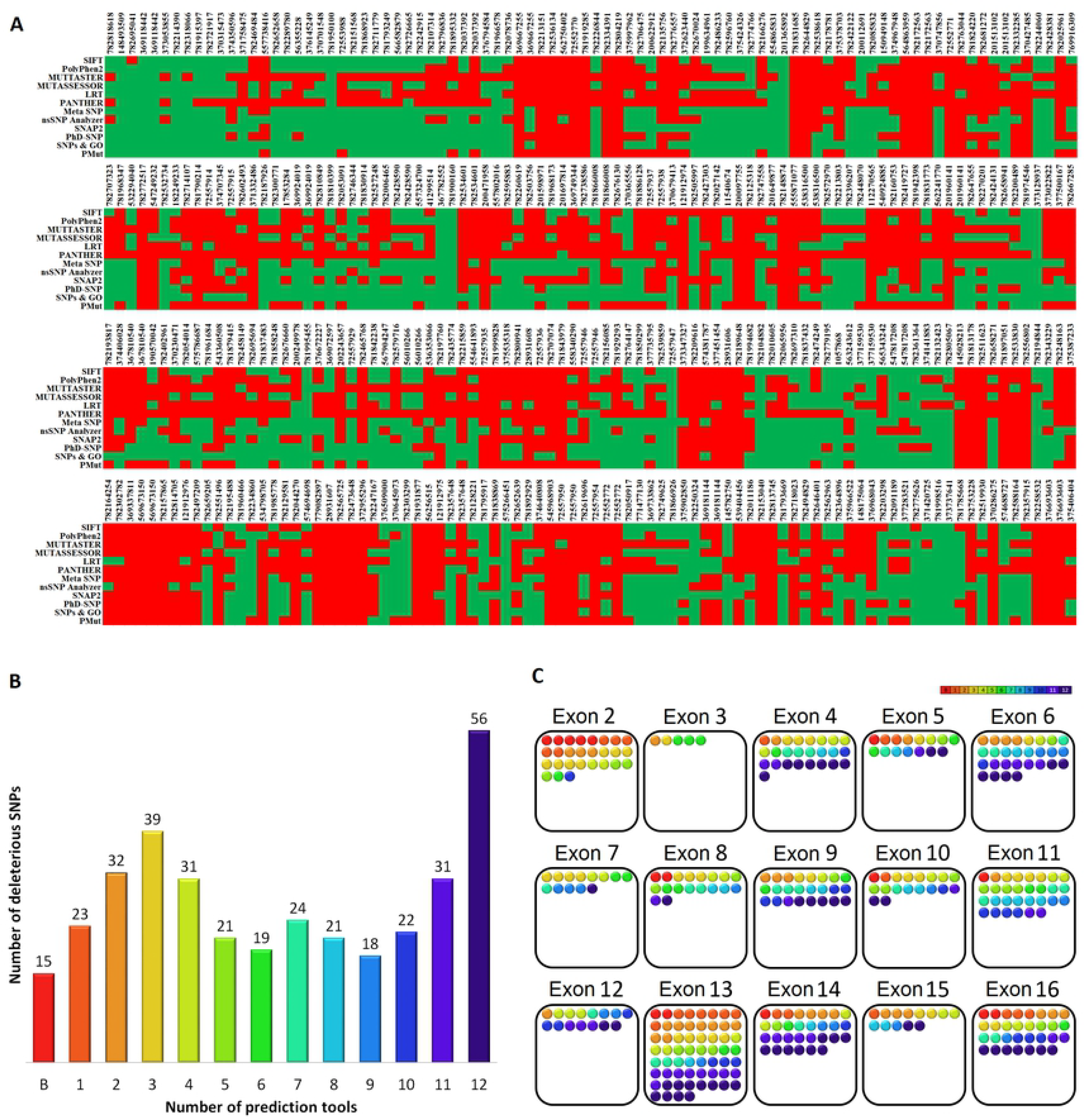
Prediction of deleterious variants using different tools. (A) Benign and deleterious effect of missense variants predicted by each tool (B) Number of deleterious variants predicted by corresponding number of tools applied. B denotes the benign SNPs (C) Distribution of benign and deleterious SNPs in each exon of POR gene.

The highly deleterious variants were subjected to evolutionary conservation analysis using ConSurf server to identify the highly conserved high risk nsSNPs (Fig.3). Six percentage of the amino acid residues in POR protein sequence had structural significance, whereas 14.8 % were functionally important. Out of 87 high risk nsSNPs, 58 had Consurf score of 8 and 9, making them highly conserved and these variants were designated as very high risk nsSNPs. Only three SNPs that changes amino acids in the positions 138, 169 and 661 of POR protein (buried region), were found to lack structural and functional conservation. At positions 177, 202, 301, 492, 517, 535, 537, 569, 570, 600 and 673, the alternate possibilities of amino acid substitutions were also found to be very high risk.

**Fig. 3:**
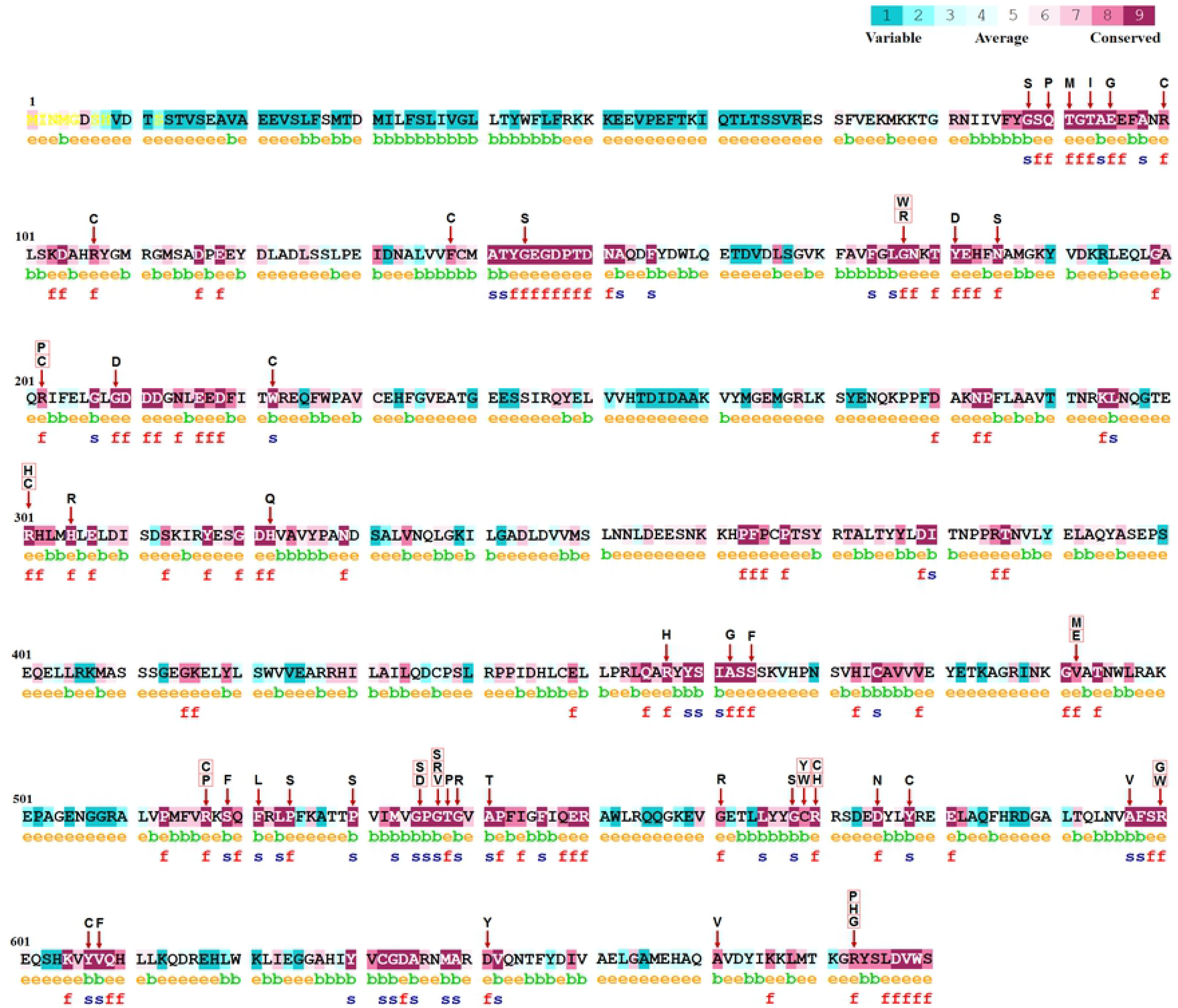
Very high risk missense variants predicted by ConSurf. The red arrow shows the change of amino acid caused by very high risk missense variants. The boxed amino acid codes represents the highly deleterious substitutions by more than one amino acid in that location due to codon change. The color code of aminoacids indicate the conservation score based on the scale at the top. e – exposed residue; b – buried residue; s – highly conserved structural residue; f – highly conserved functional residue.

A total of 58 very high risk nsSNPs were then subjected to MutPred2 analysis to elucidate the structural and functional consequences in molecular mechanisms. MutPred2 score > 0.75 were considered pathogenic and the corresponding disrupted molecular mechanisms with probability score > 0.35 and p-value < 0.01 are given in Table 1. The complete list of very high risk nsSNPs along with their MutPred2 results are given as supplementary data (**Table S3**). The very high risk variants were found to alter the structure and function of POR protein significantly.

**Table 1:** MutPred results of very high risk nsSNPs predicted with threshold MutPred2 score > 0.75, disrupted molecular mechanism probability > 0.35 and p-value < 0.01.

| Variant | MutPred2 Score | Molecular mechanisms disrupted | Probability | P-value |
| --- | --- | --- | --- | --- |
| Q90P | 0.943 | Loss of Catalytic site at Q90 | 0.56 | 6.80E-05 |
| T93I | 0.795 | Altered Metal binding | 0.41 | 1.10E-03 |
| E95G | 0.892 | Altered Metal binding | 0.62 | 1.90E-03 |
| R100C | 0.775 | Altered Metal binding | 0.44 | 8.00E-04 |
|  |  | Loss of Relative solvent accessibility | 0.38 | 1.10E-03 |
| G177R | 0.902 | Altered Ordered interface | 0.42 | 4.60E-04 |
| G177W | 0.908 | Altered Ordered interface | 0.4 | 8.60E-04 |
| Y181D | 0.939 | Altered Metal binding | 0.45 | 6.30E-03 |
|  |  | Altered Ordered interface | 0.35 | 5.00E-03 |
| G209D | 0.851 | Altered Metal binding | 0.43 | 9.70E-05 |
| G535D | 0.901 | Altered Metal binding | 0.39 | 2.30E-04 |
|  |  | Gain of Allosteric site at G535 | 0.36 | 2.50E-04 |
|  |  | Loss of Catalytic site at P536 | 0.35 | 7.40E-04 |
| G537R | 0.924 | Gain of Allosteric site at G537 | 0.55 | 4.90E-06 |
|  |  | Altered Metal binding | 0.41 | 9.70E-06 |
| G537V | 0.927 | Loss of Catalytic site at P536 | 0.37 | 6.10E-04 |
|  |  | Altered Metal binding | 0.35 | 7.80E-05 |
| T538P | 0.913 | Altered Metal binding | 0.35 | 1.20E-03 |
| G539R | 0.863 | Gain of Allosteric site at G539 | 0.4 | 1.50E-04 |
|  |  | Altered Metal binding | 0.38 | 2.90E-05 |
| C569W | 0.937 | Altered Metal binding | 0.43 | 5.60E-03 |
|  |  | Altered Ordered interface | 0.36 | 3.60E-03 |
| Y578C | 0.795 | Altered Metal binding | 0.51 | 4.70E-03 |
| Y607C | 0.789 | Altered Metal binding | 0.58 | 3.10E-03 |
| D641Y | 0.906 | Altered Metal binding | 0.47 | 4.50E-03 |
|  |  | Loss of Relative solvent accessibility | 0.39 | 9.80E-04 |

The very high risk variants were predicted to affect the ligand binding property of POR protein by altering the catalytic site amino acids. The residues involved in ligand binding are represented in Fig. 4, the purple arrows indicate the very high risk variants and red arrow indicates the affected catalytic sites. The positions Q90 and P536 are involved in FMN and NADP binding, respectively, which may be affected by the polymorphisms Q90P, G535D and G537V.

**Fig.4:**
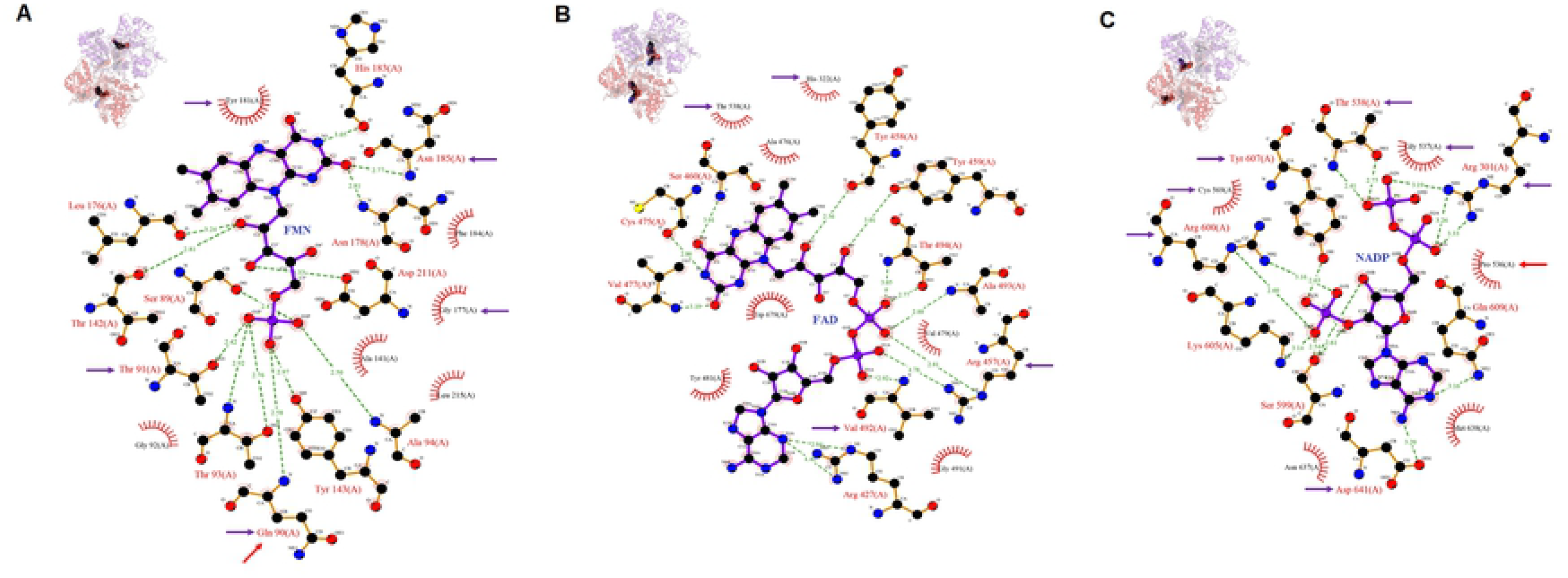
Very high risk variants involved in binding of ligands, FMN (A), FAD (B) and NADP (C). The top protein tertiary structure shows the ligand bound to it. Green dotted lines indicate hydrogen bonds and red ray like structures indicate non – bonded interactions. Purple and red arrows represent the positions of very high risk missense variants and most significantly affected catalytic residues, respectively.

The non-coding SNPs with minor allele frequency (MAF) > 0.01 in POR gene were retrieved using HaploReg v4, which gives MAF of four different populations, African, American, Asian and European. Among 1046 non-coding SNPs present in POR gene, 133 SNPs were found to have MAF > 0.01 in all the four populations. Significant difference in MAF was observed among Africans compared to other populations. HaploReg v4 also revealed the non-coding variants located in the regulatory regions associated with binding of transcription factors (TFs) and enhancers (Fig.5a).

**Fig.5:**
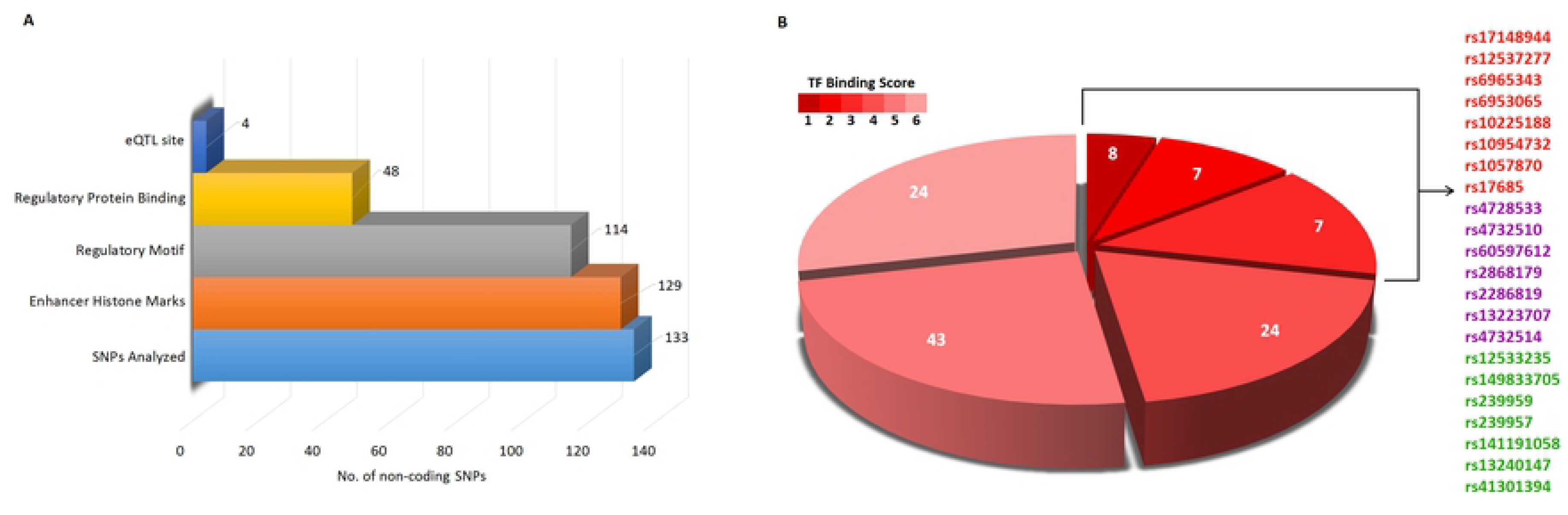
Very high risk non-coding variants identified by HaploReg v4 and RegulomeDB. (A) Functional annotation of non-coding variants with MAF > 0.01. (B) Distribution of non-coding SNPs based on RegulomeDB score. The SNPs with score 1, 2 and 3 are likely to affect TF binding and alter gene expression and the colors red, purple and green represents the scores of SNPs, respectively.

These variants were then analyzed with RegulomeDB tool to predict the influence of polymorphism with TF binding and underlying gene expression. Out of 133 SNPs with MAF > 0.01, TF binding data was obtained for 114 SNPs, in which 22 SNPs were found to likely affect TF binding with RegulomeDB score ≤ 3. The eight non-coding SNPs with least score, rs17148944, rs12537277, rs6965343, rs6953065, rs10225188, rs10954732, rs1057870 and rs17685 were considered as very high risk.

The very high risk SNPs in miRNA target site were identified using three tools, PolymiRTs v3.0, miRNASNP2 and MirSNP. Three out of 13 SNPs in miRNA target regions were predicted by all the three tools, which were considered as very high risk (Table 2). These SNPs were found to break the miRNA target site in most cases, preventing gene expression regulation by miRNAs. The miRNA, hsa-miR-185-3p was found to target the region where two very high risk SNPs, rs112507039 and rs184275273 are located. In addition, seven SNPs, rs13921, rs17685, rs41302345, rs41302348, rs72557960, rs72557961 and rs55909219 were predicted to affect miRNA target site by two of the three tools used.

**Table 2:**
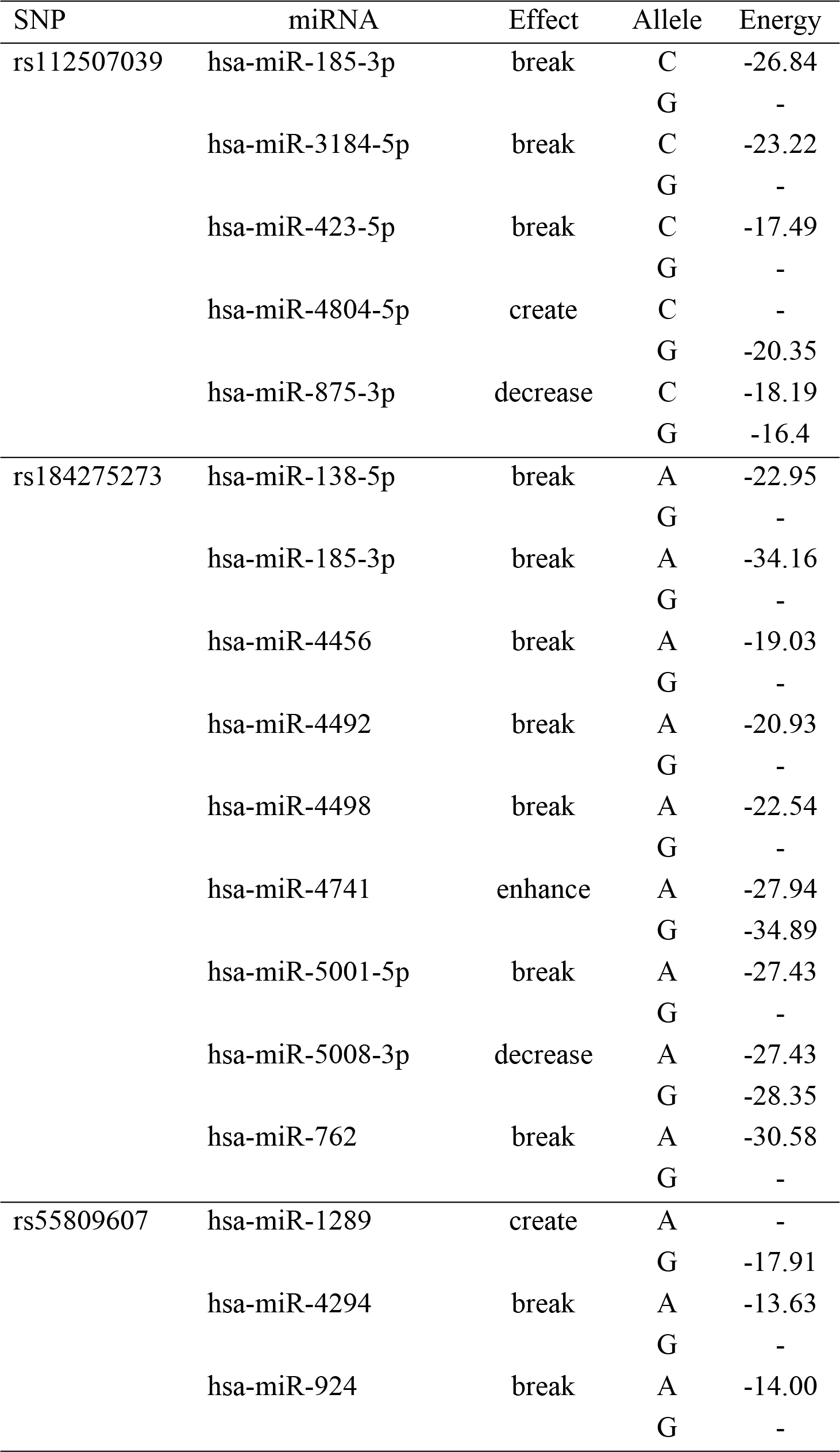
Very high risk SNPs affecting miRNA target regions.

| SNP | miRNA | Effect | Allele | Energy |
| --- | --- | --- | --- | --- |
| rs112507039 | hsa-miR-185-3p | break | C | -26.84 |
|  |  |  | G | - |
|  | hsa-miR-3184-5p | break | C | -23.22 |
|  |  |  | G | - |
|  | hsa-miR-423-5p | break | C | -17.49 |
|  |  |  | G | - |
|  | hsa-miR-4804-5p | create | C | - |
|  |  |  | G | -20.35 |
|  | hsa-miR-875-3p | decrease | C | -18.19 |
|  |  |  | G | -16.4 |
| rs184275273 | hsa-miR-138-5p | break | A | -22.95 |
|  |  |  | G | - |
|  | hsa-miR-185-3p | break | A | -34.16 |
|  |  |  | G | - |
|  | hsa-miR-4456 | break | A | -19.03 |
|  |  |  | G | - |
|  | hsa-miR-4492 | break | A | -20.93 |
|  |  |  | G | - |
|  | hsa-miR-4498 | break | A | -22.54 |
|  |  |  | G | - |
|  | hsa-miR-4741 | enhance | A | -27.94 |
|  |  |  | G | -34.89 |
|  | hsa-miR-5001-5p | break | A | -27.43 |
|  |  |  | G | - |
|  | hsa-miR-5008-3p | decrease | A | -27.43 |
|  |  |  | G | -28.35 |
|  | hsa-miR-762 | break | A | -30.58 |
|  |  |  | G | - |
| rs55809607 | hsa-miR-1289 | create | A | - |
|  |  |  | G | -17.91 |
|  | hsa-miR-4294 | break | A | -13.63 |
|  |  |  | G | - |
|  | hsa-miR-924 | break | A | -14.00 |
|  |  |  | G | - |

## DISCUSSION

Cytochrome P450 oxidoreductase is involved in the metabolism of drugs and xenobiotics through the CYP enzymes. Knock out of POR gene in mice resulted in diminished hepatic drug metabolism and embryonic lethality [11]. The highly polymorphic characteristic feature of POR gene plays major role in the differential enzyme activity and underlying phenotypic changes in drug metabolism. Several missense and other types of mutations and polymorphisms were studied in POR gene in association with diseases and drug metabolism. In this study, the very high risk SNPs in POR gene, which may significantly affect the enzyme phenotype were predicted using robust computational tools.

Among 352 missense variants analyzed, 58 were found to be very high risk and their phenotypic effect was predicted using MutPred2 tool. With the threshold value of MutPred2 score > 0.75, disrupted molecular mechanism probability > 0.35 and p-value < 0.01, it was found that three very high risk missense variants at positions 90, 535 and 537 may affect the catalytic site of FMN and NADP at positions Q90 and P536, respectively. The variants in these positions are rs782536134 (Q90P), rs782302782 (G535S), rs369337811 (G535D), rs569673150 (G537S/R) and rs782157865 (G537V).

Fluck et al., studied the impact of several missense mutations and polymorphisms in POR gene on its enzyme activity. The very high risk variant Y181D was found to eliminate 99% of POR activity due to loss of hydrophobicity in FMN binding domain. Very high risk variants in FAD binding sites R457 (rs28931608) and V492 (rs377451454, rs28931606) resulted in complete loss of POR activity. The common missense variant A503V which is present in the unstructured loop was not identified as very high risk with the computational tools used in this study, however further functional characterizations are required in confirming the phenotypic effect of this variant. The very high risk variants G539R, C569Y and V608F were found to result in reduced binding of NADPH and subsequent decrease in POR activity but not completed loss [12].

Out of 133 non-coding SNPs with MAF > 1%, eight were considered very high risk based on HaploReg v4 and RegulomeDB analysis. The variants rs17148944 and rs1057870 were found to contribute in the estimation of warfarin maintenance dose in addition to other potential SNPs in *POR, CYP2CP, VKORC1* and *CYP4F2* [4]. The very high risk variant rs6965343 was recently found to be associated with adverse effects of anti-hepatitis B drug, Pradefovir, in Chinese population [14].

A total of 13 SNPs were identified in miRNA target regions of POR gene and among these three SNPs, rs112507039, rs184275273 and rs55809607 were found to be very high risk which are predicted by all three prediction tools. Further functional studies are essential to explore the role of these miRNAs and miRSNPs in POR activity.

## CONCLUSION

POR is a highly polymorphic protein with significant role in pharmacogenetics and other physiological functions. Among 4,601 POR variants of different functional classes, 58 missense variants, eight non-coding SNPs and three SNPs in miRNA target regions were identified as very high risk SNPs in POR gene. Population based analysis of these variants in POR will reveal their significance in the metabolism of drugs and xenobiotics.

## Supplementary data

**S1 Table. List of genetic variants in cytochrome P450 oxidoreductase**

**S2 Table. Data on functional analysis of nsSNPs in POR gene using 12 different prediction tools.**

**S3 Table. Complete MutPred2 results of all very high risk missense variants.**

## REFERENCES

1. Yamano S, Aoyama T, McBride OW, Hardwick JP, Gelboin HV, Gonzalez FJ. Human NADPH-P450 oxidoreductase: complementary DNA cloning, sequence and vaccinia virus-mediated expression and localization of the CYPOR gene to chromosome 7. Mol Pharmacol. 1989; 36:83–88.

2. Wang M, Roberts DL, Paschke R, Shea TM, Masters BS, Kim JJ. Three-dimensional structure of NADPH-cytochrome P450 reductase: prototype for FMN- and FAD-containing enzymes. Proc Natl Acad Sci U S A. 1997; 94:8411–8416.

3. Scott RR, Gomes LG, Huang NW, Van Vliet G, Miller WL. Apparent manifesting heterozygosity in P450 oxidoreductase deficiency and its effect on coexisting 21-hydroxylase deficiency. J Clin Endocrinol Metab. 2007; 92:2318–2322.

4. Zhang X, Li L, Ding X, Kaminsky LS. Identification of cytochrome P450 oxidoreductase gene variants that are significantly associated with the interindividual variations in warfarin maintenance dose. Drug Metab Dispos 2011; 39:1433–1439.

5. Hart SN, Wang S, Nakamoto K, Wesselman C, Li Y, Zhong XB. Genetic polymorphisms in cytochrome P450 oxidoreductase influence microsomal P450-catalyzed drug metabolism. Pharmacogenet Genomics 2008; 18:11–24.

6. Zhang HF, Li ZH, Liu JY, Liu TT, Wang P, Fang Y, et al. Correlation of cytochrome P450 oxidoreductase expression with the expression of 10 isoforms of cytochrome P450 in human liver. Drug Metab Dispos 2016; 44:1193–1200.

7. Xiao X, Ma G, Li S, Wang M, Liu N, Ma L, et al. Functional POR A503V isassociated with the risk of bladder cancer in a Chinese population. Sci Rep 2015; 5:11751.

8. Huang N, Agrawal V, Giacomini KM, Miller WL. Genetics of P450 oxidoreductase: sequence variation in 842 individuals of four ethnicities and activities of 15 missense mutations. Proc Natl Acad Sci U S A. 2008; 105:1733–1738.

9. Pandey AV, Sproll P. Pharmacogenomics of human P450 oxidoreductase. Front Pharmacol 2014; 5:103.

10. Lv J, Hu L, Zhuo W, Zhang C, Zhou H, Fan L. Effects of the selected cytochrome P450 oxidoreductase genetic polymorphisms on cytochrome P450 2B6 activity as measured by bupropion hydroxylation. Pharmacogenet Genomics 2016; 26:80–87.

11. Shen A.L., O’Leary K.A., Kasper C.B. Association of multiple developmental defects and embryonic lethality with loss of microsomal NADPH-cytochrome P450 oxidoreductase. J. Biol. Chem. 2002; 277: 6536–6541.

12. Flück CE, Nicolo C, Pandey AV. Clinical, structural and functional implications of mutations and polymorphisms in human NADPH P450 oxidoreductase. Fundam Clin Pharmacol. 2007;21(4):399–410.

13. Ding Y, Zhang H, Li X, Li C, Chen G, Chen H, Wu M, Niu J. Safety, pharmacokinetics and pharmacogenetics of a single ascending dose of pradefovir, a novel liver-targeting, anti-hepatitis B virus drug, in healthy Chinese subjects. Hepatol Int. 2017;11(4):390–400.

